# ADAMIXTURE: Adaptive First-Order Optimization for Biobank-Scale Genetic Clustering

**DOI:** 10.64898/2026.02.13.700171

**Authors:** Joan Saurina-i-Ricos, Daniel Mas Montserrat, Alexander G. Ioannidis

## Abstract

**Motivation:** Estimating genetic clusters from sequencing data is a fundamental task in population and medical genetics, enabling demographic inference and adjustment for population structure in association studies. ADMIXTURE, a widely used model-based clustering method, employs an accelerated Expectation–Maximization (EM) algorithm to infer population parameters; however, its computational demands scale poorly, limiting its usefulness for modern biobank-sized datasets. While recent EM acceleration strategies employing second-order quasi-Newton schemes preserve accuracy, they remain computationally intensive. Conversely, EM-free approaches that prioritize speed often compromise solution quality.

**Results:** We introduce ADAMIXTURE, a novel optimization framework that integrates the EM algorithm with Adaptive Moment Estimation (Adam). Unlike traditional acceleration methods, ADAMIXTURE utilizes first-order gradients with adaptive learning rates derived from raw and squared moments to approximate curvature information, bypassing the computational overhead of Hessian approximations. This approach surpasses the convergence efficiency of second-order methods while maintaining the low computational complexity of first-order updates. Across simulated and large-scale empirical datasets, ADAMIXTURE demonstrates substantial reductions in wall-clock runtime and enhanced scalability compared to state-of-the-art methods, while maintaining comparable or improved inference accuracy. Its GPU implementation runs in under 2 hours on half a million samples and variants, a two order of magnitude speedup over current state-of-the-art.

**Availability and implementation:** Source code is available at: https://github.com/AI-sandbox/ADAMIXTURE.

## 1 Introduction

The rapid expansion of population-scale biobanks and sequencing initiatives has revolutionized the genomic prediction of human traits and disease risk. However, the predictive accuracy of these models is often linked to genetic ancestry, rendering the precise characterization of population structure a prerequisite for robust analysis [16]. For decades, unsupervised ancestry inference has served as a cornerstone for correcting stratification in genome-wide association studies and reconstructing demographic histories. Yet, as datasets scale to millions of individuals and genomic variants [4], foundational algorithms increasingly face computationally prohibitive bottlenecks. Overcoming these scalability constraints is paramount for analyzing diverse biobanks and ensuring that the promise of precision medicine is equitably realized across all populations.

The standard computational framework for this task, established by `STRUCTURE` [19], models individuals as mosaics of fractional ancestries derived from *K* ancestral populations. These populations are inferred via unsupervised learning in a high-dimensional space defined by variant frequencies, allowing for the characterization of continuous genetic variation without reliance on subjective labels. The model operates on single nucleotide polymorphisms (SNPs), exploiting the fact that allele frequency distributions diverge among populations due to distinct demographic processes, including migration, genetic drift, and founder effects.

While the Bayesian framework introduced by `STRUCTURE` was seminal, its reliance on Markov Chain Monte Carlo (MCMC) sampling imposed severe computational costs on large datasets. To mitigate this, `ADMIXTURE` [1] reformulated the problem within a maximum likelihood framework, employing block relaxation and Expectation-Maximization (EM) algorithms to accelerate inference. This shift significantly extended the applicability of genetic ancestry estimation and clustering. Nevertheless, with the rapid growth of large-scale genomic cohorts, the computational demands of `ADMIXTURE` have likewise become prohibitive, rendering it intractable for biobank-scale analysis.

In response, recent years have seen the emergence of diverse acceleration strategies, broadly categorized into likelihood-free and likelihood-based approaches. Likelihood-free methods include matrix factorization such as `sNMF` [9], which has been superseded in biobank-scale applications by the more scalable alternating least squares (ALS) scheme in `SCOPE` [5]. Conversely, likelihood-based accelerators focus on optimizing the EM procedure itself; for instance, `fastmixture` [21] combines mini-batch EM optimization with ALS-driven initialization. Concurrently, alternative paradigms have been adapted to this domain, such as `Archetypal Analysis` [10] and deep learning approaches like `Neural Admixture` [7], which respectively recasts genetic ancestry inference and clustering as an autoencoder task. While these innovations have improved scalability, they often introduce significant trade-offs between computational speed and statistical rigor.

Addressing this dichotomy, we introduce **ADAMIXTURE**, a novel optimization framework that accelerates the EM algorithm by integrating it with Adaptive Moment Estimation (Adam) [13]. Unlike prior acceleration attempts, our method leverages first-order *pseudo-gradients* to drive robust parameter updates within a single EM step. ADAMIXTURE is available in both CPU (Central Processing Unit) and GPU (Graphics Processing Unit) implementations. While the CPU version ensures broad accessibility, the GPU variant leverages highly parallel computation to boost scalability and throughput. The result is a powerful tool achieving state-of-the-art performance on biobank-scale datasets.

## 2 Subjects and methods

### 2.1 Model formulation

This work builds upon the probabilistic framework introduced in `STRUCTURE`, which describes the relationship between an observed genotype matrix, *G*, and two latent parameter matrices: the individual admixture proportions, *Q*, and the ancestral allele frequencies, *F* .

The model takes as input the *M × N* genotype matrix *G*, where each entry *G*_*ji*_ ∈ {0, 1, 2}contains the count of alternate alleles for SNP *j* in individual *i*, across *M* SNPs and *N* individuals. The parameter matrices are:

- **Admixture Proportions (Q):** A *K × N* matrix where *q*_*ki*_ is the proportion of individual *i*’s ancestry from ancestral population *k*, such that 0 *≤ q*_*ki*_ *≤* 1 and 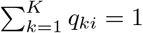.
- **Allele Frequencies (F):** An *M× K* matrix where *f*_*jk*_ is the frequency of the alternate allele at SNP *j* in ancestral population *k*, such that 0 *≤ f*_*jk*_ *≤* 1.

The model connects these by defining an expected individual-specific allele frequency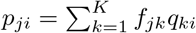, which represents the weighted average of the ancestral frequencies. The observed genotype *G*_*ji*_ is then modeled as a random draw from a binomial distribution, *G*_*ji*_ *∼*Binomial(2, *p*_*ji*_). The primary goal is to infer the latent matrices *Q* and *F* from the observed genotypes *G*.

### 2.2 Memory optimization

Because each observed genotype entry *G*_*ji*_ takes only one of three discrete states, the dataset can be compactly encoded using low-precision formats. We utilize an 8-bit encoding for CPU computations and a 2-bit encoding for GPU computations, converting to a 64-bit format strictly for the data subset actively being processed. This approach reduces the data transfer overhead between the disk, CPU, and GPU while minimizing the overall memory footprint. A description of the memory requirements is detailed in the supplementary material (Supplementary Note S1).

### 2.3 Parameter estimation

The parameters of the model are estimated by maximizing the likelihood of the observed genotype data *G*. This is equivalent to minimizing the negative log-likelihood function. Assuming each genotype *G*_*ji*_ is an independent draw from a binomial distribution, the negative log-likelihood is given by:

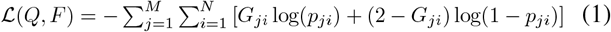

Subject to the parameter constraints, the optimization problem becomes:

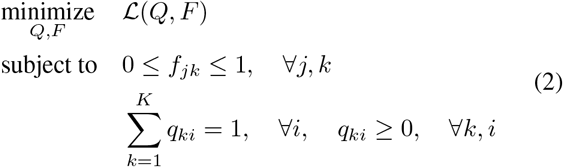

### 2.4 Expectation–Maximization

We employ the EM procedure introduced in `FRAPPÉ` [23] and later adopted by `ADMIXTURE`, which iteratively updates the ancestry proportions *Q* and allele frequencies *F* .

In the Expectation step, we compute 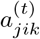 and 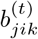, the posterior probabilities that an observed alternate or reference allele, respectively, originated from ancestral population *k*:

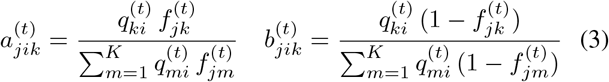

In the Maximization step, the model parameters are updated by maximizing the expected complete-data log-likelihood with respect to the current posterior assignments:

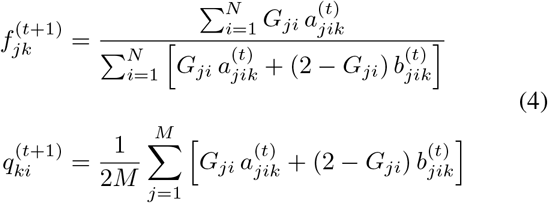

This procedure alternates between computing expected ancestry assignments and updating parameter estimates until convergence.

### 2.5 Initialization

The choice of starting values for ancestry proportions *Q* and allele frequencies *F* is critical for ensuring convergence in EM-based admixture models. Naive random initialization often results in entrapment in suboptimal local maxima, a problem that is further exacerbated by accelerated algorithms.

While `Neural Admixture` [7] employs soft clustering on data projected via Principal Component Analysis (PCA) to provide informative initialization points, it struggles to scale to large numbers of ancestral populations or very large sample sizes. To address this, we adopt and refine the initialization strategy employed by `SCOPE` [5] and `fastmixture` [21], incorporating both an advanced spectral decomposition and an Alternating Least Squares (ALS) factorization framework. Specifically, we implement randomized Singular Value Decomposition (SVD) with Dynamic Shifts [8], a technique that utilizes dynamically shifted power iterations to refine the approximation of leading singular vectors.

The resulting low-rank structure approximates the centered genotype matrix, which serves as the target for the ALS factorization. This procedure jointly estimates the starting *Q* and *F* matrices prior to EM optimization. While `SCOPE` and `fastmixture` rely on naive box-clipping to project parameters in each iteration to satisfy the constraints in Eq. (2), such an approach—which simply truncates out-of-bound parameters to the nearest edge of the [0, 1] interval—fails to redistribute the remaining variables to account for the underlying dependency. This abrupt truncation displaces the solution from the true gradient path, inducing oscillatory convergence patterns or even preventing convergence altogether at a large number of populations or under conditions of high ancestral correlation.

To preserve the true gradient and strictly satisfy the Karush-Kuhn-Tucker (KKT) optimality conditions across the entire feasible region, we formulate the row-wise ALS updates as Bounded Variable Least Squares (BVLS) problems [22]. We solve these efficiently via a 3-state variant of the Block Principal Pivoting (BPP) algorithm [12], which dynamically manages free variables alongside lower and upper active bounds to enforce the 0 ≤ *x* ≤ 1 constraints. After each update, to strictly satisfy the constraints formulated in Eq. (2) and avoid numerical instabilities, *F* and *Q* are capped slightly inside the boundaries and the rows of *Q* are *L*_1_-normalized, mapping admixture proportions onto the interior of the *K*-dimensional probability simplex. A detailed description of the initialization is provided in the supplementary material (Supplementary Note S2).

In order to demonstrate the empirical advantages of our initialization strategy, we evaluated its performance across a range of ancestral populations using 100,000 samples and 100,000 SNPs from the UK Biobank. Across all tested scenarios, our approach yields a significantly higher initial value of the objective function *ℒ* (*Q, F*) compared to `fastmixture`, leading to faster convergence (Figure 3) and improved final objective values. Furthermore, our initialization method consistently captures the expected scaling behavior with an increasing number of populations leading to higher log-likelihoods, as the larger parameter space accommodates finer ancestral resolution. Detailed comparisons of the initial and resulting log-likelihood values are provided in the supplementary material (Supplementary Note S3).

### 2.6 Adam-EM

Standard EM algorithms exhibit linear convergence rates, which become prohibitively slow on biobank-scale datasets. Established fixed-point acceleration schemes seek to bypass this bottleneck, but they often introduce substantial per-iteration overhead. For instance, the Squared Iterative Method (SQUAREM) [24] and the second-order quasi-Newton algorithms employed by `ADMIXTURE` and `fastmixture` [26] typically necessitate two EM evaluations per parameter update to formulate their extrapolations or approximate the Hessian. Similarly, Anderson acceleration [25] requires storing a history of previous residuals and solving a least-squares problem at each iteration. These requirements effectively double the computational cost per step or introduce severe memory and processing bottlenecks in high-dimensional settings.

#### Algorithm 1

Adam-EM

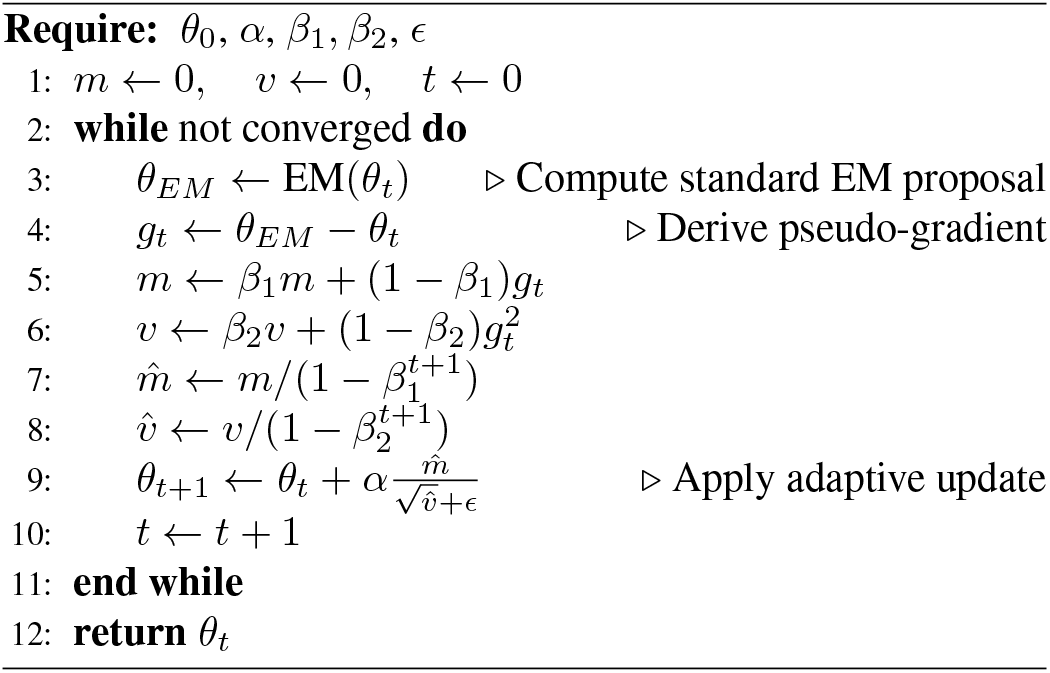

To overcome this limitation, we introduce **Adam-EM**, an optimization method that integrates the stability of EM with the adaptive dynamics of the Adam optimizer. This approach uses first-order information, unlike second-order methods. The EM-induced parameter shift can be interpreted as a *pseudogradient*—a proxy for the steepest ascent direction on the likelihood manifold [14]. This interpretation is grounded in the fact that the EM update represents the true gradient transformed by a positive-definite matrix, effectively acting as a first-order method operating on a locally reshaped likelihood surface [20]. Feeding this *pseudo-gradient* into Adam allows adaptive moment estimation to normalize updates across parameters with different sensitivities. This effectively approximates curvature and injects momentum without the computational overhead of Hessian construction. The detailed pseudocode for this general optimization strategy is presented in Algorithm 1. In this generic formulation, the stopping criterion is application-specific, typically based on thresholding relative parameter changes or monitoring the target objective function.

Beyond genetic clustering and ancestry inference, our framework shows potential applicability to the broad class of latent variable models where EM updates are computationally expensive and exact gradients are intractable—such as probabilistic matrix factorization and topic modeling. Its underlying mechanism, which decouples the search direction (provided by EM) from the step size dynamics (managed by Adam), provides a theoretical foundation for its use in other settings characterized by high dimensionality and ill-conditioned parameter spaces. While further validation is needed beyond genetics, the method offers a scalable alternative that combines first-order stability with convergence speed that matches or exceeds second-order methods.

### 2.7 Optimization

We apply the Adam-EM acceleration scheme to simultaneously optimize the admixture proportions *Q* and allele frequencies *F* . In each iteration, we utilize the standard EM update rules derived in Eqs. (3)–(4) to compute the *pseudo-gradients*, following the logic described in Algorithm 1. After the adaptive step, parameters are projected to strictly satisfy the constraints formulated in Eq. (2) by applying box-clipping to the [0, 1] interval for both matrices, followed by *L*_1_-normalization of the rows of *Q*. Moment buffers (*m, v*) are not reset upon parameter clipping, as potential boundary chattering is effectively mitigated by our step-size adaptation mechanism.

To promote numerical stability and efficient convergence, we incorporate a dynamic learning rate schedule that monitors the objective function *ℒ* (*Q, F*) defined in Eq. (1). Computing the log-likelihood is costly, and storing biobank-scale matrices for rollbacks incurs substantial memory overhead, so we save parameters only when the objective improves. Furthermore, to reduce floating-point error accumulation—particularly in momentum-based schemes [11]—the objective is evaluated every *c* iterations. If no improvement occurs over *ρ* consecutive checks, the learning rate is reduced by a factor *γ ∈* (0, 1). Optimization halts safely once the learning rate reaches a minimum *α*_min_. This strategy allows occasional likelihood-worsening steps within the *c*-iteration window but serves as a robust step-size heuristic, with subsequent iterations at the reduced rate refining the solution. The complete procedure is detailed in Algorithm 2, and all hyperparameters are provided in the supplementary material (Supplementary Note S4).

#### Algorithm 2

ADAMIXTURE

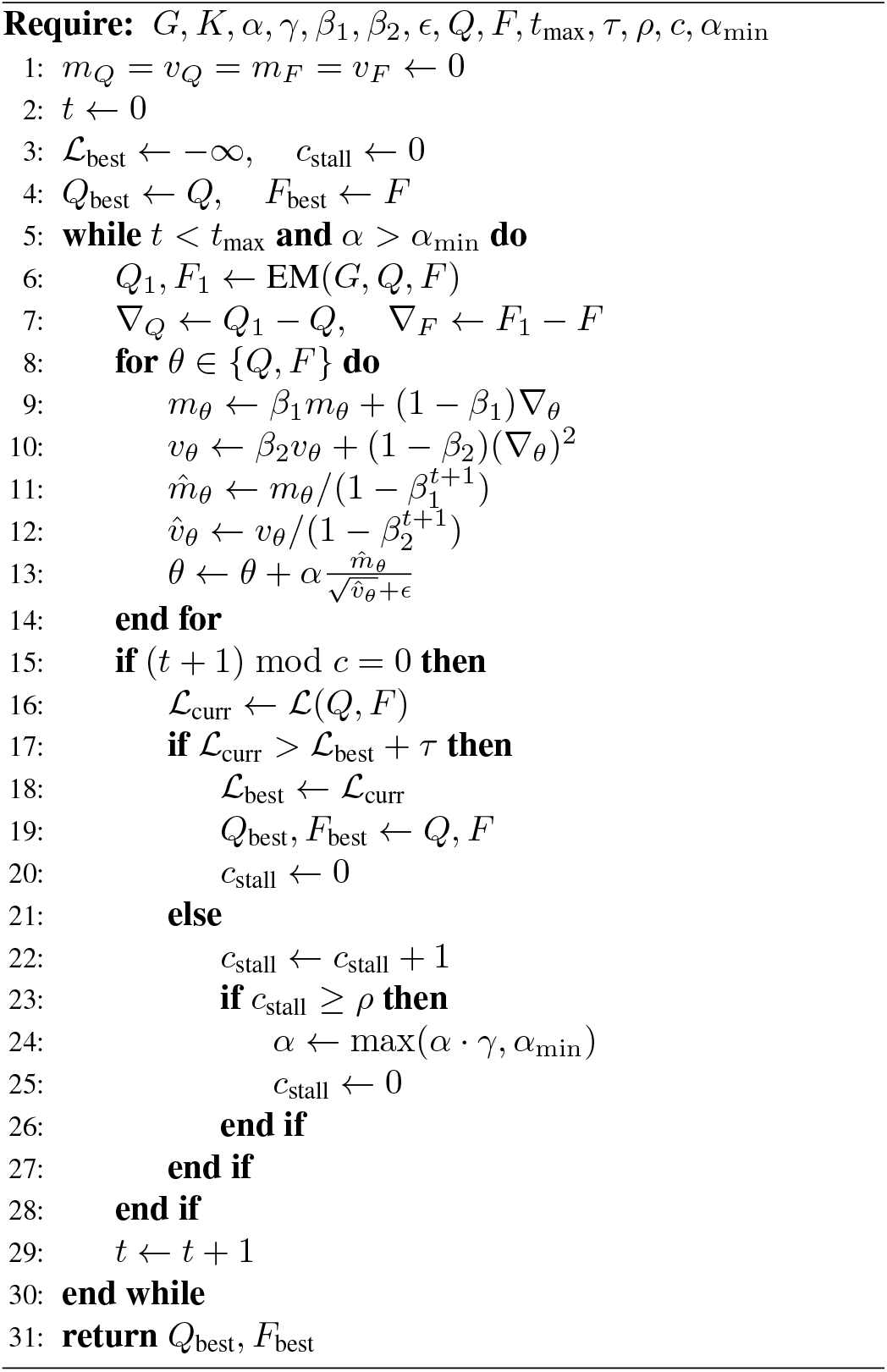

## 3 Benchmarking

### 3.1 Assessment of results

We assess method performance through a strategy encompassing quantitative goodness-of-fit metrics, reproducibility analysis, and qualitative inspection of population structure. Given the unsupervised paradigm of genetic ancestry inference, where absolute ground truth is absent in empirical datasets, we rely on the negative log-likelihood from Eq. (1) as the primary objective function. This metric quantifies the statistical fidelity of the parameter estimates and the model’s capacity to explain the observed genetic variation.

To evaluate the robustness of the inferred ancestry components across independent initializations, we measure cluster stability by computing the mean pairwise Pearson correlation and the Frobenius distance between *Q* matrices over 5 independent runs per configuration. This provides a reproducibility check, ensuring that the results are consistent and not artifacts of local minima.

Complementing statistical fit and stability, we benchmark computational efficiency by recording wall-clock runtime across varying dataset dimensions to evaluate scalability and qualitatively validate the results by visualizing the *Q* matrices using `pong` [2]. All results from empirical and simulated datasets are publicly available on [figshare](https://doi.org/10.6084/m9.figshare.31065292).

### 3.2 Simulated datasets

To enable direct comparison with state-of-the-art acceleration methods, we use the simulation benchmarks introduced by `fastmixture` [21], generated via coalescent simulations with a constant effective population size of 10,000 to model neutral evolution.

We consider four demographic scenarios (A–D). **Scenarios A, B**, and **C** represent discrete populations with constant sizes, while **Scenario D** models a complex American-Admixture event [3]. Sample sizes are 1,000 for Scenarios A, B, and D, and 1,600 for Scenario C. After filtering for minor allele frequency (MAF) *>* 0.05, the resulting datasets contain between 500,000 and 700,000 SNPs.

### 3.3 Empirical datasets

To evaluate ancestry inference under realistic conditions, we selected four genomic cohorts representing distinct regimes of sample size, SNP density, and population structure. This diverse set provides a robust framework for assessing both scalability and accuracy, capturing patterns from fine-scale stratification to global population divergence.

The **UK Biobank (UKB)** [4] serves as our primary benchmark for biobank-scale inference. For the central accuracy and runtime comparisons, we curated a standard subset comprising 100,000 individuals and 100,000 SNPs, obtained via random subsampling and MAF filtering. To systematically evaluate algorithmic scalability across different data dimensions, we also generated auxiliary subsets with sample sizes ranging from 50,000 to 500,000 individuals and SNP counts from 100,000 to 500,000. Predominantly of European ancestry, this cohort challenges both scale and fine-structure resolution, reflecting modern Genome-Wide Association Study (GWAS) contexts.

The **Domestic Dog** dataset [18] includes 1,355 samples from 166 breeds genotyped at 150,131 SNPs. Originating from lineages across all inhabited continents, the cohort exhibits strong genetic drift and sharp population boundaries due to breed isolation. It provides a rigorous test of model robustness and adaptability, particularly for inferring high numbers of ancestral populations.

The merged dataset of the **Human Genome Diversity Project (HGDP)** and **1000 Genomes Project** comprises 4,091 individuals from globally distributed populations. After MAF filtering, the panel includes*∼* 500,000 autosomal SNPs, with sex chromosomes excluded. Compared to UK Biobank, this panel captures substantial worldwide genetic diversity and pronounced population structure, making it a complementary benchmark for evaluating performance under complex, multi-ancestry scenarios [6, 15].

### 3.4 Experimental setup

All experiments were conducted on the UCSC Genomics Institute Phoenix cluster. Each computational run was allocated 64 logical cores on an AMD EPYC 7713 processor. As `Neural Admixture` and the GPU implementation of ADAMIXTURE are the only methods capable of leveraging hardware acceleration, they were each provisioned with a single NVIDIA A100 GPU (80 GB VRAM); all other methods were restricted to CPU execution.

We benchmark ADAMIXTURE against `ADMIXTURE`, `Neural Admixture`, `fastmixture`, and `SCOPE`, with details on software versions, libraries, and execution parameters provided in the supplementary material (Supplementary Note S5).

## 4 Results

### 4.1 Performance on Simulated Benchmarks

Quantitative benchmarking on simulated data confirms that ADAMIXTURE consistently recovers the global maximum with numerical precision and stability comparable to or exceeding that of `ADMIXTURE` and `fastmixture`. While all three likelihood-based methods converge to a very similar global mode, our method achieves marginally higher absolute loglikelihood values across all simulated datasets, demonstrating comparable convergence tightness to the baselines (Table 1).

**Table 1:** Log-likelihood measures on Simulated Datasets. Each dataset was run with the specified number of clusters: Dataset A (*K* = 3), B (*K* = 4), C (*K* = 5), D (*K* = 3). For each method, the mean of 5 independent runs is reported, with the standard deviation shown in parentheses.

| Method | A | B | C | D |
| --- | --- | --- | --- | --- |
| ADAMIXTURE | -637,729,492.3 (0.1) | -629,360,597.6 (0.0) | -1,006,294,343.7 (0.1) | -429,286,152.3 (0.1) |
| fastmixture | -637,729,494.0 (0.7) | -629,360,599.1 (0.3) | -1,006,294,347.8 (0.5) | -429,286,152.5 (0.1) |
| Neural Admixture | -637,765,240.0 ($3.6 \times 10^4$) | -630,345,588.3 ($8.4 \times 10^5$) | -1,007,821,071.2 ($6.6 \times 10^3$) | -429,332,568.9 ($2.9 \times 10^4$) |
| ADMIXTURE | -637,729,492.4 (0.5) | -629,360,598.5 (1.6) | -1,006,294,347.6 (4.1) | -429,286,155.6 (4.1) |
| SCOPE | -637,846,223.5 (10.2) | -629,538,208.7 (38.1) | -1,006,604,603.8 (244.3) | -429,584,894.3 (84.4) |

In contrast, approximate methods fail to reach this optimality. `SCOPE` consistently produces suboptimal fits, falling short of the likelihood quality achieved by EM-based approaches. Similarly, `Neural Admixture` exhibits high variance across replicates, yielding unreliable parameter estimates.

### 4.2 Performance on Empirical Benchmarks

Evaluation on large-scale empirical cohorts further validates the efficacy of our framework. As detailed in Table 2, ADAMIXTURE attains the maximum log-likelihood across all evaluated datasets. Crucially, our method demonstrates superior numerical stability, maintaining low standard deviations across replicates, whereas competing accelerated methods frequently exhibit substantial variance. Across all cohorts, `Neural Admixture` and `SCOPE` systematically fail to reach the optimal mode, yielding model fits with likelihoods orders of magnitude worse than those of state-of-the-art methods.

**Table 2:** Log-likelihood measures on Empirical Datasets. Each dataset was run with the specified number of clusters: UKB (*K* = 5), DOGS (*K* = 20), HGDP (*K* = 8). For each method, the mean of 5 independent runs is reported, with the standard deviation shown in parentheses.

| Method | UKB | Dogs | HGDP |
| --- | --- | --- | --- |
| ADAMIXTURE | -5,324,953,196.1 (21.4) | -194,648,323.0 ($1.6 \times 10^4$) | -1,131,620,423.1 (0.2) |
| fastmixture | -5,325,159,277.5 ($3.2 \times 10^5$) | -194,651,453.1 ($1.1 \times 10^4$) | -1,131,620,449.9 (6.5) |
| Neural Admixture | -5,334,181,181.0 ($1.3 \times 10^5$) | -195,266,696.7 ($8.4 \times 10^4$) | -1,132,157,396.4 ($3.5 \times 10^5$) |
| ADMIXTURE | -5,324,953,302.9 (32.0) | -194,749,866.8 ($6.4 \times 10^4$) | -1,131,908,125.6 ($5.0 \times 10^5$) |
| SCOPE | -5,340,156,808.4 ($3.5 \times 10^6$) | -195,985,480.7 (63.4) | -1,134,190,154.7 ($2.8 \times 10^3$) |

While `fastmixture` and classical `ADMIXTURE` generally converge toward near-optimal solutions, our method consistently outperforms both baselines in precision and convergence robustness. The practical impact of this improved robustness is illustrated in Figure 1. Qualitative inspection shows that the ancestry proportions inferred by ADAMIXTURE are visually indistinguible from those recovered by `fastmixture` and classical `ADMIXTURE`, demonstrating that the proposed acceleration strategy preserves fine-scale population structure.

**Figure 1:**
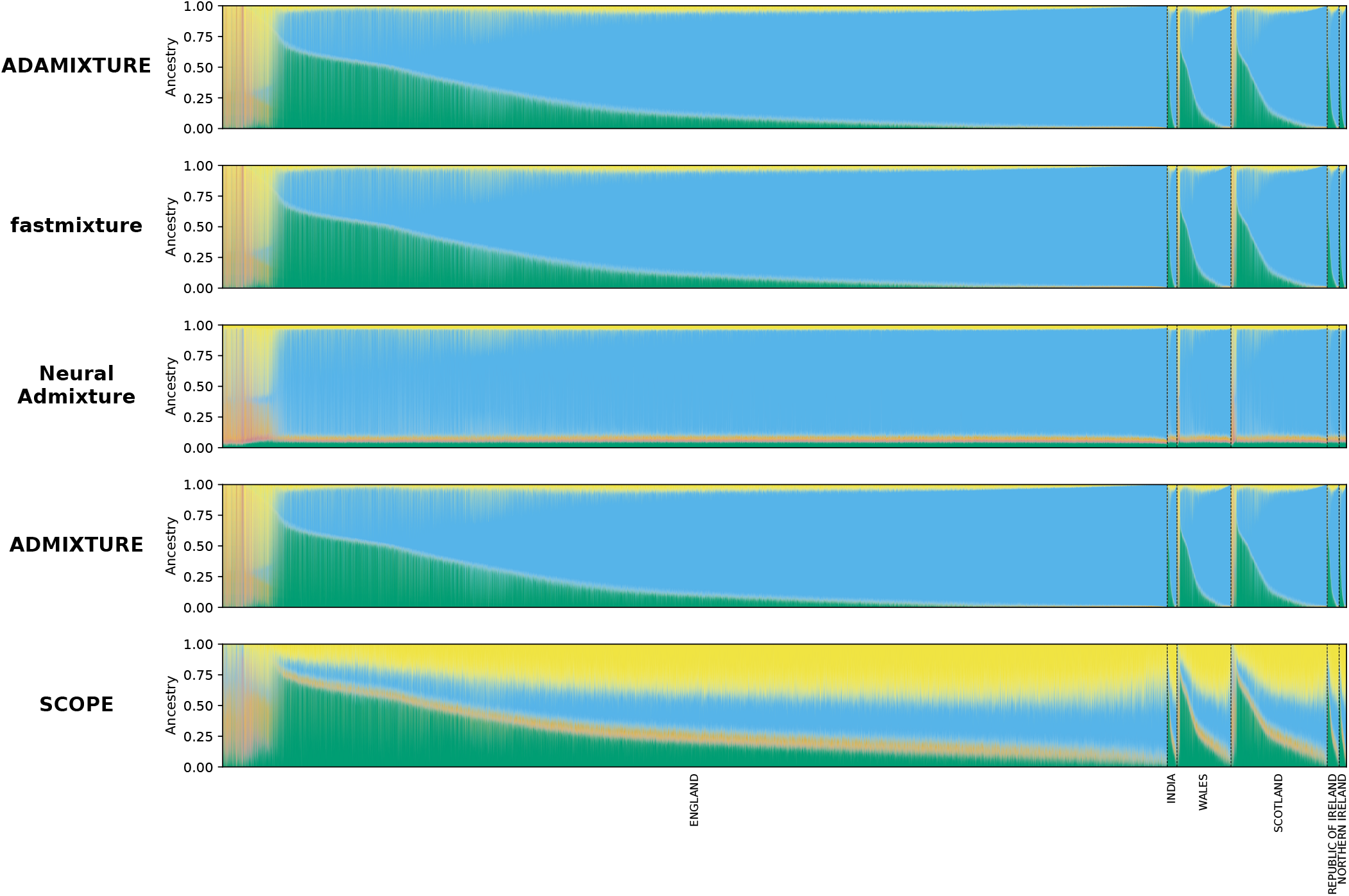
Visualization of the estimated ancestry proportions for the UK Biobank dataset (*K* = 5) obtained using the different benchmarked methods. Each individual is represented by a thin vertical line partitioned into colored segments that denote their estimated fractional membership in each of the *K* ancestral clusters. For each method, the displayed results correspond to the run achieving the highest log-likelihood among 5 independent runs. Labels correspond to self-reported location of birth. Only regions with a sufficient number of individuals to ensure clear visualization are shown; these include England, India, Wales, Scotland, the Republic of Ireland, and Northern Ireland.

Conversely, likelihood-free accelerators introduce significant artifacts. Under default settings, `Neural Admixture` over-smooths estimates, assigning similar ancestry proportions to distinct sub-populations—a behavior reminiscent of mode collapse or aggressive regularization. Although neural methods are highly sensitive to parameter tuning and may improve with extensive optimization, these default-setting inaccuracies corroborate prior reports of suboptimal neural-network fits [21]. On the other hand, `SCOPE` yields noisy clusters that obscure underlying genetic signals. Notably, `SCOPE`’s solutions closely resemble those obtained using only our spectral initialization, without EM refinement. This indicates that its ALS-based strategy may stagnate at a superficial local minimum, explaining the fast runtime but substantially lower solution quality.

### 4.3 Quantitative Stability and Precision

To evaluate the reliability of inferred population structures, we analyzed cluster stability and reconstruction error across independent runs. As detailed in the supplementary material (Supplementary Note S6), ADAMIXTURE demonstrates exceptional robustness, maintaining a mean pairwise Pearson correlation of *≈*1.0 across replicates for all datasets.

In terms of numerical precision, our method achieves exceptionally low Frobenius distances, outperforming likelihood-free accelerators in most scenarios while matching or exceeding the precision of likelihood-based implementations. These findings confirm that the proposed acceleration strategy does not compromise model fidelity, demonstrating robustness to the stochasticity introduced during the initialization phase.

### 4.4 Runtime and Accuracy Trade-off

The runtime efficiency of ADAMIXTURE was systematically benchmarked against competing methods across all evaluated datasets, achieving substantial acceleration over classical methods (Figure 2). The most substantial performance gain is observed when compared to the standard `ADMIXTURE` algorithm on the largest empirical dataset (100,000 samples). While `ADMIXTURE` requires over 57 hours to reach convergence, our baseline CPU implementation completes the inference in approximately 45 minutes. Crucially, the introduction of hardware acceleration via ADAMIXTURE (GPU) further reduces this runtime to a mere 5 minutes, achieving a staggering 680*×* overall speedup. This effectively transforms the analysis of biobank-scale cohorts from a multi-day task into a sub-hour routine.

**Figure 2:**
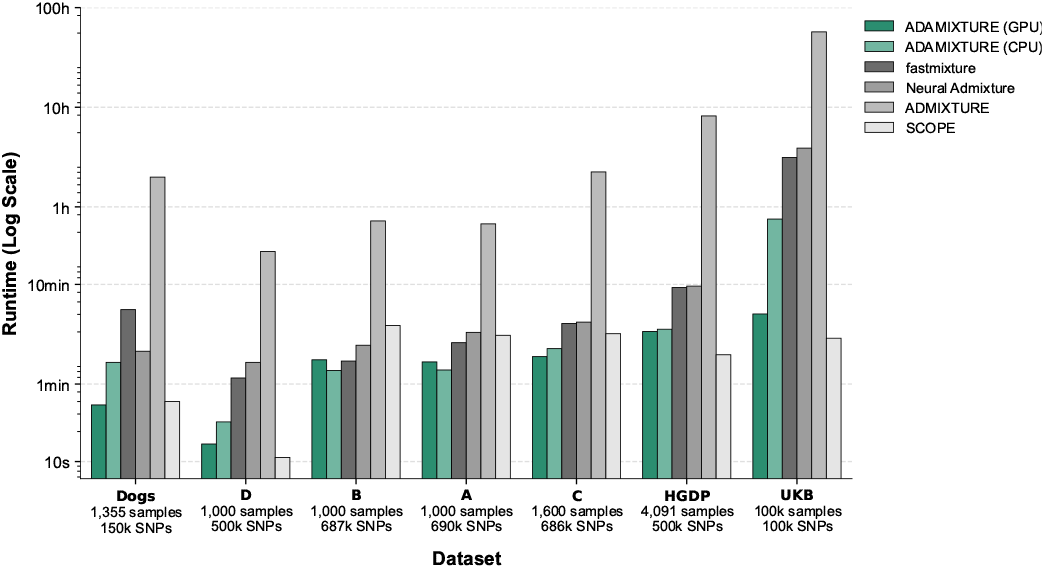
Comparison of runtime performance across synthetic (A-D) and empirical (Dogs, HGDP, UKB) datasets. The y-axis is shown on a logarithmic scale to accommodate the extreme variance in execution times. Results represent the mean of 5 independent runs.

On smaller cohorts, the CPU–GPU performance gap is narrow because initialization dominates execution time in low-complexity regimes. However, as dataset dimensions scale up, the GPU-accelerated Adam-EM optimization becomes the defining factor, as demonstrated in the subsequent sections.

Furthermore, ADAMIXTURE consistently outperforms other modern accelerated frameworks. On average across all evaluated cohorts, it achieves a 3.5*×* speedup over `fastmixture` and a 4*×* speedup over `Neural Admixture`. A nuanced profile emerges for `SCOPE`, which exhibits the lowest absolute wall-clock times on datasets with moderate feature dimensionality (*<* 500, 000 SNPs). Nevertheless, both ADAMIXTURE variants prove more computationally efficient on high-dimensional datasets (synthetic sets A–C). Importantly, as established in our prior analyses, `SCOPE`’s speed advantage in some scenarios is consistently achieved at the expense of solution quality.

### 4.5 Scalability Analysis

To ensure a rigorous assessment of computational efficiency, we restrict our scalability benchmarks to ADAMIXTURE and `fastmixture`. Heuristic approximations such as `SCOPE` and `Neural Admixture` are excluded from this direct comparison because, as demonstrated in Tables 1 and 2, they consistently fail to converge to the global maximum; their lower runtimes derive in part from premature termination at suboptimal optima rather than fully from algorithmic efficiency. Moreover, `ADMIXTURE` is omitted due to its prohibitive computational cost. As shown in Figure 2, its execution on a UK Biobank subset (*K* = 5) already approaches 60 hours, rendering it intractable for the configurations explored in this evaluation.

We evaluate how the number of ancestral populations affects processing time using a fixed subset of the UK Biobank (100, 000 samples, 100, 000 SNPs). For this benchmark, the maximum iteration limit for both methods was increased to 10, 000, as the default limits were insufficient to guarantee convergence for high values of *K*. Additionally, some hyperparameters for ADAMIXTURE were adjusted, with details provided in the supplementary material (Supplementary Note S4).

As illustrated in Figure 3, ADAMIXTURE (CPU) exhibits profoundly superior performance scaling compared to `fastmixture`. The execution of `fastmixture` becomes impractical as the number of ancestral populations increases, with runs for *K ≥*25 failing to complete within a 10-day limit. In contrast, our CPU version demonstrates near-logarithmic execution growth across the entire evaluated range up to *K* = 50.

**Figure 3:**
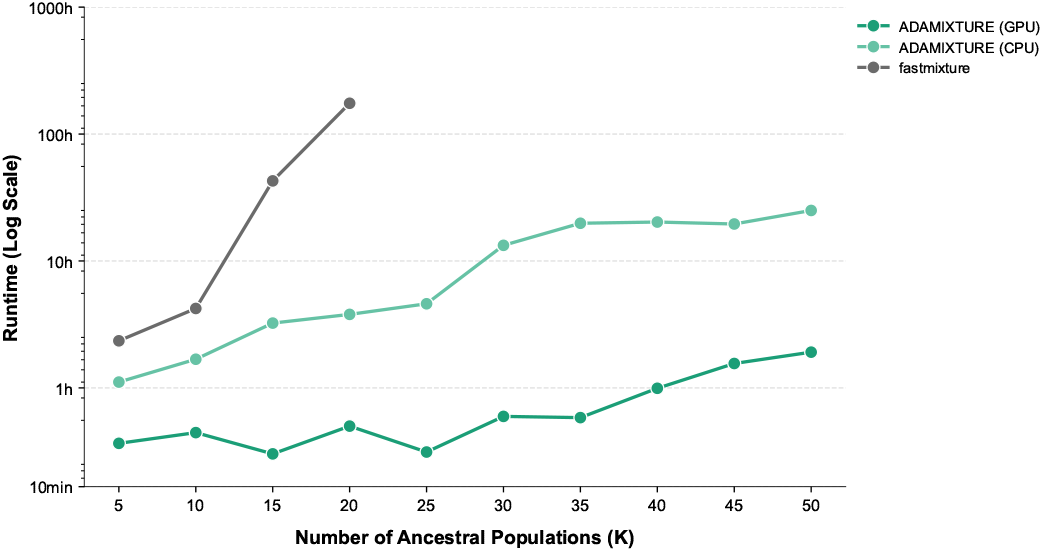
Runtime scalability with respect to the Number of Ancestral Populations (*K*) on a subset of the UK Biobank dataset (100, 000 samples, 100, 000 SNPs). Results represent the mean of 5 independent runs. Data points for `fastmixture` at *K ≥*25 are omitted as they exceeded a 10-day runtime limit.

This efficiency gain is intrinsic to our Adam-EM formulation. By eliminating the necessity for the traditional two-step EM procedure, our method effectively halves the processing overhead associated with updating the population matrices at each step. Importantly, this drastic reduction in per-iteration cost is not offset by a need for more iterations. As shown in the supplementary material (Supplementary Note S3), ADAMIXTURE converges in a comparable or lower number of iterations while simultaneously reaching better solutions.

Additionally, the GPU implementation of ADAMIXTURE further reduces execution time by an order of magnitude. Even at high complexity (*K* = 50), convergence occurs in approximately 2 hours, making previously prohibitive biobank-scale clustering tasks routine.

Beyond the number of ancestral populations, we also assess data scaling by varying the number of samples and SNPs while maintaining a fixed number of populations. As illustrated in Figure 4, ADAMIXTURE (CPU) consistently outperforms `fastmixture` across all evaluated configurations. Moreover, ADAMIXTURE (GPU) achieves a paradigm shift in execution speed, dramatically accelerating convergence. At the maximum evaluated scale, the GPU version converges in under 2 hours, whereas the baseline `fastmixture` requires nearly 100 hours. This performance gap highlights ADAMIXTURE’s exceptional suitability for massive cohorts, offering a fast, robust solution that scales naturally without compromising statistical rigor.

**Figure 4:**
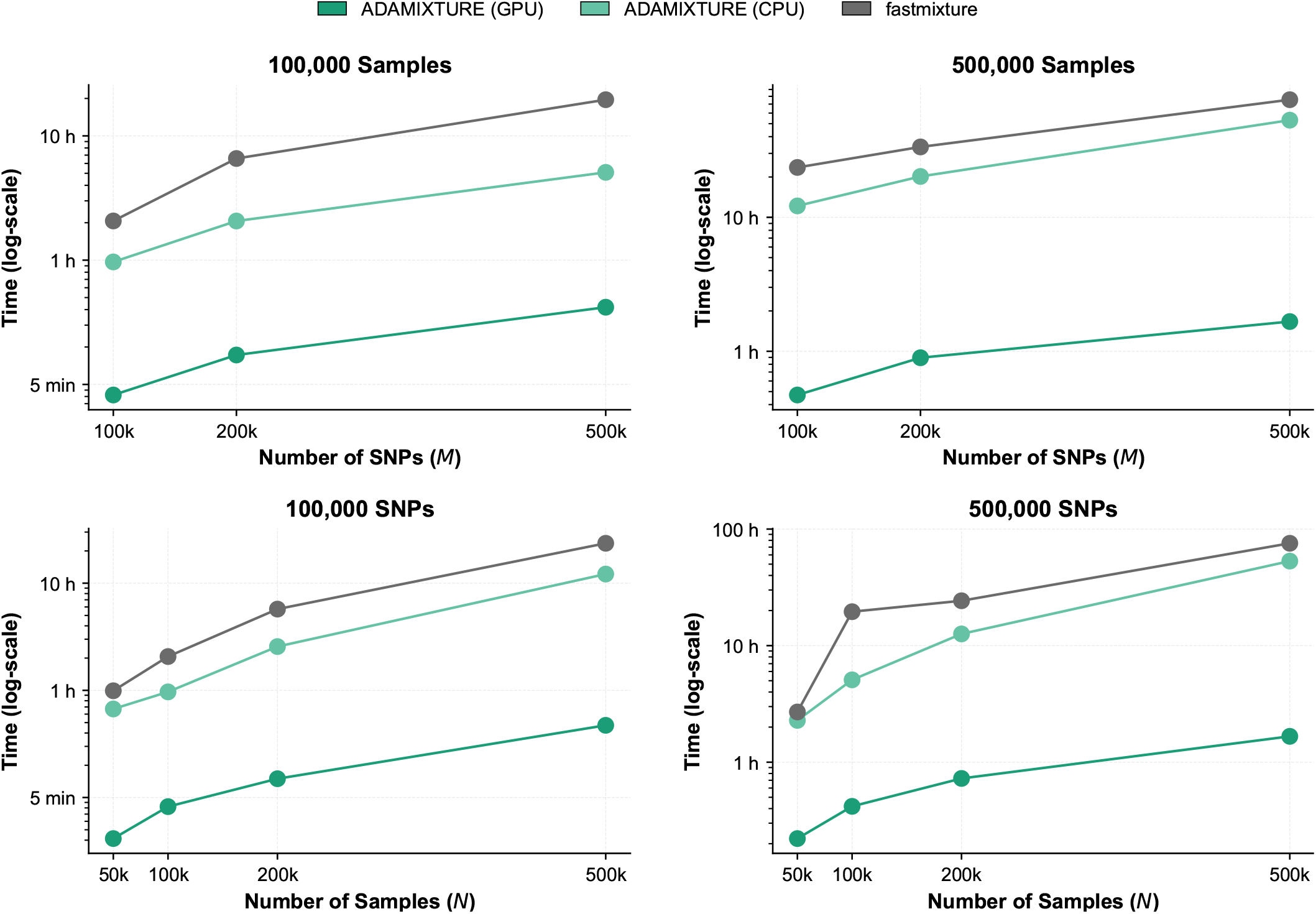
Scalability with respect to dataset dimensions. The plot illustrates runtime growth on the UK Biobank dataset as a function of the number of SNPs (top panels) and sample size (bottom panels), with the number of populations fixed at *K* = 5. Results represent the mean of 5 independent runs.

Taken together, these results demonstrate that the primary processing bottleneck for standard CPU-based EM methods lies in the number of estimated populations rather than the raw dimensions of the genotype data. While scaling samples and markers yields a predictable, manageable increase in execution time, increasing the number of latent populations severely slows traditional algorithms. It is precisely in overcoming this fundamental limitation that the GPU-accelerated version of ADAMIXTURE demonstrates its greatest advantage, seamlessly absorbing the workload of high-dimensional clustering.

## 5 Discussion

This study demonstrates that integrating first-order adaptive optimization with the EM framework effectively resolves the scalability bottlenecks currently plaguing genetic clustering. Our findings indicate that the heavy computational expense of second-order updates is redundant in high-dimensional genomic settings, where adaptive momentum can achieve comparable convergence tightness with significantly lower overhead.

Our results underscore that the precision of the initialization phase is as critical as the optimization itself. While previous accelerators relied on heuristic clipping that often derailed the initial gradient path, the shift toward an exact BVLS formulation ensures that the optimization consistently begins within the basin of the global maximum.

Recognizing that the analysis of full-scale biobanks still poses a non-trivial time investment on standard CPU hardware, our framework is natively designed to harness massive hardware parallelism. Through its GPU-accelerated implementation, we effectively bridge the gap between statistical guarantees and ultra-fast execution. While prior attempts like `Neural Admixture` utilized GPUs but sacrificed likelihood maximization for speed, our approach achieves the optimal model fit in a fraction of the time. This delivers an additional order-of-magnitude reduction in runtime, compressing biobank-scale analyses that previously required days or weeks into the order of an hour, even at extreme levels of complexity.

Looking forward, on the computational front, extending our implementation to multi-GPU architectures and mixed-precision computing presents a straightforward pathway to further accelerate inference for unprecedented cohort sizes, although special care must be taken to mitigate numerical imprecision introduced by distributed gradient reductions and reduced-precision formats. In terms of downstream applications, a critical next step will be evaluating the impact of these highly resolved *Q*-matrices on population stratification correction in complex trait GWAS, as well as improving the cross-population portability of Polygenic Risk Scores (PRS). Furthermore, adapting the Adam-EM framework to operate on haplotype clusters, which have recently demonstrated superior accuracy in ancestry estimation compared to unlinked SNPs [17], or extending it to accelerate local ancestry inference, represents a highly relevant methodological extension. Finally, exploring the integration of this optimizer with models designed for low-coverage sequencing data or ancient DNA could also broaden its utility.

## 6 Author contributions

J.S.R., A.G.I., and D.M.M. designed the research. J.S.R. wrote the software. J.S.R., D.M.M., and A.G.I. interpreted the results. J.S.R., D.M.M., and A.G.I. wrote the manuscript.

## 7 Acknowledgements

This work uses data provided by patients and collected by the NHS as part of their care and support. We thank the UK Biobank and its participants and the NHS. This research has been conducted using the UK Biobank Resource under Application Number 24983.

## Notes

https://github.com/AI-sandbox/ADAMIXTURE

https://doi.org/10.6084/m9.figshare.31065292

### Competing Interest Statement

The authors have declared no competing interest.

### Summary of Updates

A GPU implementation is added that increases the speed of the method when GPUs are available. Additional clarifications are also introduced.

