## Supplementary Material for "ADAMIXTURE: Adaptive First-Order Optimization for Biobank-Scale Genetic Clustering"

Joan Saurina-i-Ricos, Daniel Mas Montserrat, Alexander G. Ioannidis

#### SUPPLEMENTARY NOTE S1: PEAK MEMORY COMPLEXITY ANALYSIS

To provide complete transparency regarding the computational resources required by ADAMIXTURE, we detail the exact theoretical bounds for peak memory usage during the EM-Adam optimization phase. It is critical to emphasize that in biobank-scale genetic clustering, the dominant memory footprint is dictated by the input genotype matrix $G$. Because $G$ is common to all clustering algorithms, this baseline spatial complexity represents a fundamental lower bound for any implementation. The model parameters and optimizer states are natively stored in double precision (`float64`, representing 8 bytes per value). Let $M$ be the number of SNPs, $N$ the number of samples, $K$ the number of ancestral populations, and $C$ the batch chunk size.

**1. CPU Implementation (RAM):** To optimize memory access speeds on the CPU, the genotype matrix $G$ is expanded into an 8-bit continuous array (storing 1 genotype per byte), which strictly requires $MN$ bytes. The optimizer maintains several state matrices to compute the Adam updates: 5 matrices of size $M \times K$ ($P, m_P, v_P, P_1, P_{\text{best}}$) requiring $40MK$ bytes; 6 matrices of size $N \times K$ ($Q, m_Q, v_Q, Q_1, T, Q_{\text{best}}$) requiring $48NK$ bytes; and 1 vector of size $N$ ($q_{\text{bat}}$) requiring $8N$ bytes. Combined with the unpacked genotype matrix ($MN$ bytes), the exact theoretical peak RAM (in Gigabytes) is formulated as:

$$\text{RAM}_{\text{CPU}} = \frac{40MK + 48NK + 8N + MN}{1024^3} \tag{1}$$

**2. GPU Implementation (VRAM):** To leverage hardware acceleration while strictly managing VRAM constraints, the GPU implementation processes the data in batches of size $C$ and retains the genotype matrix $G$ in its 2-bit packed PLINK format (4 genotypes per byte), requiring $\frac{MN}{4}$ bytes. The device memory footprint consists of: 6 matrices of size $M \times K$ ($P, m_P, v_P, A_{\text{accum}}, B_{\text{accum}}, P_{\text{EM}}$) totaling $48MK$ bytes; 5 matrices of size $N \times K$ ($Q, m_Q, v_Q, T_{\text{accum}}, Q_{\text{EM}}$) totaling $40NK$ bytes; 6 temporary batch buffers of size $C \times N$ (`float64`) and 1 unpacked memory buffer totaling $52CN$ bytes. The exact theoretical peak VRAM (in Gigabytes) is formulated as:

$$\text{VRAM}_{\text{GPU}} = \frac{48MK + 40NK + 52CN + \frac{MN}{4}}{1024^3} \tag{2}$$

**Numerical Examples ($K = 5$):** To illustrate the scalability of ADAMIXTURE, we evaluate two high-dimensional scenarios:

- **Standard Biobank** ($100,000 \times 100,000$)**:** On **CPU**, the 8-bit genotype matrix occupies $\approx 9.31$ GB, with the Adam state adding only $\approx 0.04$ GB (Total: **9.35 GB**). On **GPU** ($C = 1024$), the 2-bit matrix occupies $\approx 2.33$ GB and batch buffers $\approx 4.96$ GB (Total: **7.33 GB**).

- **Extreme Scale** ($500,000 \times 500,000$)**:** On **CPU**, the 8-bit genotype matrix dominates with $\approx 232.83$ GB, while the Adam state remains minimal at $\approx 0.21$ GB (Total: **233.04 GB**). On **GPU** ($C = 1024$), the 2-bit matrix requires $\approx 58.21$ GB and the batch buffers $\approx 24.79$ GB, totaling **83.20 GB** of VRAM.

These results confirm that even at extreme scales, the optimizer's overhead is negligible ($< 0.1\%$ on CPU). Notably, the GPU VRAM requirements can be further reduced by tuning the chunk size $C$, allowing for execution on professional hardware even at half-million scales.

**Practical Overhead:** The formulas presented above represent the raw tensor allocations. In practical deployment, these estimates should be scaled by a safety margin of approximately 5% to 10% ($\sim \times 1.05$ to $\times 1.10$). This overhead accounts for the initial CUDA context reservation, PyTorch's internal execution engine structures, and the behavior of the dynamic garbage collector.

### SUPPLEMENTARY NOTE S2: ADAMIXTURE INITIALIZATION

Algorithm S1 details the overall initialization procedure. The process begins by obtaining a low-rank approximation of the centered genotype matrix via randomized Singular Value Decomposition (SVD). To ensure a biologically plausible and mathematically robust starting point for the subsequent Expectation-Maximization (EM) steps, we rely on an Alternating Least Squares (ALS) framework. Rather than applying standard unconstrained least squares followed by post-hoc boundary clipping—which ignores the covariance structure between ancestral populations and disrupts the gradient path—we formulate the row-wise updates as exact Bounded Variable Least Squares (BVLS) problems.

For each update step (e.g., updating $F$), we first compute the unconstrained target ($F_{\text{free}}$) and adjust it by the covariance matrix of the fixed variables ($A_Q$). This adjusted target ($B_F$) and the covariance matrix ($A_Q$) are then passed to Algorithm S2. Following each exact BVLS optimization, the variables are strictly constrained via the project function: allele frequencies ($F$) are bounded to $[\epsilon, 1 - \epsilon]$ to prevent numerical singularities during log-likelihood evaluation, and admixture proportions ($Q$) are additionally projected onto the strict interior of the $K$-dimensional probability simplex to sum to exactly one.

Algorithm S2 serves as the core solver for the BVLS formulation, employing a 3-state Block Principal Pivoting (BPP) method. Unlike traditional active-set methods that exchange a single variable per iteration, the 3-state BPP approach partitions the variables into a free set ($\mathcal{F}$), a lower-bound set ($\mathcal{L}$), and an upper-bound set ($\mathcal{U}$). This allows for the simultaneous exchange of multiple variables across boundaries. At each iteration, the algorithm checks for violations of the Karush-Kuhn-Tucker (KKT) optimality conditions across the entire feasible region using a numerical tolerance ($\epsilon_{\text{bpp}}$). The algorithm guarantees convergence to the exact geometric optimum within the constraints ($0 \leq x \leq 1$) in a finite number of steps, avoiding the oscillatory convergence patterns typical of naive boundary-clipping approaches. The default hyperparameters governing this initialization procedure and the BPP solver are summarized in Table S1.

**Algorithm S1** ADAMIXTURE Initialization

**Require:** $G \in \mathbb{R}^{M \times N}$, $K$, $\delta$ , $t_{\max}$, $\epsilon$, $f$, $t_{\text{bpp}}$, $\epsilon_{\text{bpp}}$

$G_c \leftarrow \text{center}(G)$

$U_{1:K}, S_{1:K}, V_{1:K}^{\top} \leftarrow \text{randomizedSVD}(G_c, K)$

$Z \leftarrow U_{1:K} S_{1:K}$ ▷ Scaled left singular vectors ($M \times K$)

$V \leftarrow (V_{1:K}^{\top})^{\top}$ ▷ Right singular vectors ($N \times K$)

% Initial Unconstrained Estimate

$F \sim \mathcal{U}(0, 1)^{M \times K}$ ▷ Initialize allele frequencies

$F \leftarrow \text{project}(F, \epsilon)$

$I_F \leftarrow F(F^{\top} F)^{-1}$

$Q \leftarrow \frac{1}{2} V Z^{\top} I_F + \mathbf{1}_N (f^{\top} I_F)$ ▷ Initial admixture estimate

$Q \leftarrow \text{project}(Q, \epsilon)$

$Q_{\text{prev}} \leftarrow Q$

$t \leftarrow 0$

**repeat**

    % 1. Update Allele Frequencies ($F$)

    $A_Q \leftarrow Q^{\top} Q$ ▷ Covariance matrix of admixture

    $I_Q \leftarrow Q A_Q^{-1}$

    $F_{\text{free}} \leftarrow \frac{1}{2} Z V^{\top} I_Q + f(\mathbf{1}_N^{\top} I_Q)$ ▷ Unconstrained target computation

    $B_F \leftarrow F_{\text{free}} A_Q$ ▷ Covariance-adjusted target for BVLS

    $F \leftarrow \text{BPP-BVLS}(A_Q, B_F, t_{\text{bpp}}, \epsilon_{\text{bpp}})$ ▷ Exact solver under KKT conditions

    $F \leftarrow \text{project}(F, \epsilon)$

    % 2. Update Admixture Proportions ($Q$)

    $A_F \leftarrow F^{\top} F$ ▷ Covariance matrix of frequencies

    $I_F \leftarrow F A_F^{-1}$

    $Q_{\text{free}} \leftarrow \frac{1}{2} V Z^{\top} I_F + \mathbf{1}_N (f^{\top} I_F)$ ▷ Unconstrained target computation

    $B_Q \leftarrow Q_{\text{free}} A_F$ ▷ Covariance-adjusted target for BVLS

    $Q \leftarrow \text{BPP-BVLS}(A_F, B_Q, t_{\text{bpp}}, \epsilon_{\text{bpp}})$ ▷ Exact solver under KKT conditions

    $Q \leftarrow \text{project}(Q, \epsilon)$

    % 3. Evaluate Convergence

    $e \leftarrow \text{RMSE}(Q, Q_{\text{prev}})$

    $Q_{\text{prev}} \leftarrow Q$

    $t \leftarrow t + 1$

**until** $e < \delta \lor t \geq t_{\max}$

**return** $Q, F$

**Algorithm S2** 3-State Block Principal Pivoting BVLS (Single Row Update)

**Require:** $A \in \mathbb{R}^{K \times K}$ (Covariance matrix), $b \in \mathbb{R}^{K}$ (Target vector), $t_{\text{bpp}}$, $\epsilon_{\text{bpp}}$

$l \leftarrow 0, \quad u \leftarrow 1$ ▷ Fixed boundary constraints for biological parameters
% Initialize index sets and vectors
$\mathcal{F} \leftarrow \{1, 2, \ldots, K\}$ ▷ Free variable set
$\mathcal{U} \leftarrow \emptyset$ ▷ Variables at upper bound
$\mathcal{L} \leftarrow \emptyset$ ▷ Variables at lower bound
$x \leftarrow \mathbf{0}^K$ ▷ Primal variables
$y \leftarrow \mathbf{0}^K$ ▷ Dual variables (Lagrange multipliers / gradients)
**for** $t = 1$ **to** $t_{\text{bpp}}$ **do**
    % Fix bounded variables
    $x_{\mathcal{U}} \leftarrow u \cdot \mathbf{1}_{|\mathcal{U}|}$
    $x_{\mathcal{L}} \leftarrow l \cdot \mathbf{1}_{|\mathcal{L}|}$
    % Adjust target vector for the influence of bounded variables
    $b' \leftarrow b - A_{:,\mathcal{U}} x_{\mathcal{U}} - A_{:,\mathcal{L}} x_{\mathcal{L}}$
    % Solve unconstrained least squares for the free variables
    $x_{\mathcal{F}} \leftarrow A^{-1}_{\mathcal{F},\mathcal{F}} b'_{\mathcal{F}}$
    % Compute the dual variables (gradients) across all parameters
    $y \leftarrow Ax - b$
    % Identify KKT condition violations
    $\mathcal{V}_{\mathcal{F}\to\mathcal{L}} \leftarrow \{i \in \mathcal{F} \mid x_i < l - \epsilon_{\text{bpp}}\}$ ▷ Free variables violating lower bound
    $\mathcal{V}_{\mathcal{F}\to\mathcal{U}} \leftarrow \{i \in \mathcal{F} \mid x_i > u + \epsilon_{\text{bpp}}\}$ ▷ Free variables violating upper bound
    $\mathcal{V}_{\mathcal{L}\to\mathcal{F}} \leftarrow \{i \in \mathcal{L} \mid y_i < -\epsilon_{\text{bpp}}\}$ ▷ Lower bounded variables that should increase
    $\mathcal{V}_{\mathcal{U}\to\mathcal{F}} \leftarrow \{i \in \mathcal{U} \mid y_i > \epsilon_{\text{bpp}}\}$ ▷ Upper bounded variables that should decrease
    **if** $\mathcal{V}_{\mathcal{F}\to\mathcal{L}} \cup \mathcal{V}_{\mathcal{F}\to\mathcal{U}} \cup \mathcal{V}_{\mathcal{L}\to\mathcal{F}} \cup \mathcal{V}_{\mathcal{U}\to\mathcal{F}} = \emptyset$ **then**
        **break** ▷ KKT conditions strictly satisfied ($l \le x \le u$)
    **end if**
    % Full Exchange Rule: Pivot variables between sets
    $\mathcal{F} \leftarrow (\mathcal{F} \setminus (\mathcal{V}_{\mathcal{F}\to\mathcal{L}} \cup \mathcal{V}_{\mathcal{F}\to\mathcal{U}})) \cup \mathcal{V}_{\mathcal{L}\to\mathcal{F}} \cup \mathcal{V}_{\mathcal{U}\to\mathcal{F}}$
    $\mathcal{U} \leftarrow (\mathcal{U} \setminus \mathcal{V}_{\mathcal{U}\to\mathcal{F}}) \cup \mathcal{V}_{\mathcal{F}\to\mathcal{U}}$
    $\mathcal{L} \leftarrow (\mathcal{L} \setminus \mathcal{V}_{\mathcal{L}\to\mathcal{F}}) \cup \mathcal{V}_{\mathcal{F}\to\mathcal{L}}$
**end for**
**return** $x$

| Parameter | Description | Value |
|---|---|---|
| $t_{\text{max}}$ | Maximum number of Alternating Least Squares (ALS) iterations. | 1000 |
| $\delta$ | Convergence tolerance threshold for the Root Mean Square Error (RMSE) between consecutive ALS iterations. | $10^{-4}$ |
| $\epsilon$ | Biological boundary constraint to prevent numerical singularities during subsequent log-likelihood evaluations. Variables are mapped marginally inside $[0, 1]$. | $10^{-5}$ |
| $t_{\text{bpp}}$ | Maximum number of iterations for the 3-State Block Principal Pivoting (BPP) BVLS solver. | 50 |
| $\epsilon_{\text{bpp}}$ | Numerical tolerance for evaluating violations of the Karush-Kuhn-Tucker (KKT) conditions in the BVLS solver. | $10^{-8}$ |

**Table S1.** Hyperparameters used in the ADAMIXTURE initialization.

### SUPPLEMENTARY NOTE S3: INITIAL AND FINAL LOG-LIKELIHOOD COMPARISON

This section evaluates the impact of the initialization strategy on the model fit using a large-scale subset of the UK Biobank (100,000 samples and 100,000 SNPs). Table S2 compares the initial log-likelihood ($\mathcal{L}$) obtained immediately after the initialization phase for both ADAMIXTURE and `fastmixture`. Table S3 summarizes the final log-likelihood values achieved at convergence, while Table S4 details the number of iterations required to reach the convergence threshold.

| K | ADAMIXTURE | fastmixture |
|---|---|---|
| 5 | **−5,325,944,414.6** | −5,335,417,978.3 |
| 10 | **−5,318,540,175.5** | −5,340,612,838.7 |
| 15 | **−5,314,308,013.2** | −5,350,310,642.7 |
| 20 | **−5,312,098,629.6** | −5,357,428,331.7 |
| 25 | **−5,310,481,145.1** | −5,371,933,837.5 |
| 30 | **−5,309,314,201.3** | −5,381,213,214.2 |
| 35 | **−5,308,362,018.0** | −5,381,607,168.2 |
| 40 | **−5,307,390,562.8** | −5,382,445,803.7 |
| 45 | **−5,306,795,396.1** | −5,389,305,119.7 |
| 50 | **−5,306,200,355.5** | −5,400,491,746.6 |

**Table S2. Initial Log-likelihood Comparison.** Comparison of the log-likelihood objective $\mathcal{L}(Q, F)$ immediately after initialization on the UK Biobank (100,000 samples, 100,000 SNPs). Results represent the mean of 5 independent runs for each method.

| K | ADAMIXTURE | fastmixture |
|---|---|---|
| 5 | **−5,324,953,252.8** | −5,324,953,717.1 |
| 10 | **−5,316,464,645.2** | −5,316,465,603.9 |
| 15 | **−5,312,196,576.5** | −5,312,255,561.8 |
| 20 | **−5,309,356,340.5** | −5,309,489,233.1 |
| 25 | **−5,307,454,805.2** | DNF |
| 30 | **−5,305,651,711.5** | DNF |
| 35 | **−5,304,214,869.6** | DNF |
| 40 | **−5,302,743,466.1** | DNF |
| 45 | **−5,301,525,485.6** | DNF |
| 50 | **−5,300,415,332.9** | DNF |

**Table S3. Final Converged Log-likelihood.** Final log-likelihood values achieved on the UK Biobank (100,000 samples, 100,000 SNPs). Results represent the mean of 5 independent runs for each method. Runs that exceeded the maximum time limit (10 days) are marked as DNF (Did Not Finish).

| K | ADAMIXTURE | fastmixture |
|---|---|---|
| 5 | 1,140 | **740** |
| 10 | 915 | **735** |
| 15 | **1,040** | 5,530 |
| 20 | **1,125** | 8,810 |
| 25 | **1,055** | DNF |
| 30 | **2,210** | DNF |
| 35 | **3,425** | DNF |
| 40 | **3,150** | DNF |
| 45 | **2,980** | DNF |
| 50 | **4,595** | DNF |

**Table S4. Iterations to Convergence.** Number of iterations required to reach the convergence threshold on the UK Biobank (100,000 samples, 100,000 SNPs). Results represent the mean of 5 independent runs. Runs that exceeded the maximum time limit (10 days) are marked as DNF (Did Not Finish).

### SUPPLEMENTARY NOTE S4: ADAMIXTURE OPTIMIZATION HYPERPARAMETERS

To ensure the reproducibility of the experiments described in the main text, Table S5 details the default hyperparameter configuration used by ADAMIXTURE.

| Parameter | Description | Value |
|---|---|---|
| $\alpha$ | Initial learning rate. | 0.005 |
| $\beta_1$ | Adam 1st moment decay. | 0.80 |
| $\beta_2$ | Adam 2nd moment decay. | 0.88 |
| $\epsilon$ | Stability constant. | $10^{-8}$ |
| $\gamma$ | Decay factor. | 0.5 |
| $\alpha_{\text{min}}$ | Minimum learning rate. | $10^{-4}$ |
| $t_{\text{max}}$ | Maximum iterations. | 1500 |
| $\tau$ | Log-likelihood tolerance. | 0.1 |
| $\rho$ | Learning rate decay patience. | 3 |
| $c$ | Log-likelihood check frequency. | 5 |

**Table S5. Default hyperparameters used for ADAMIXTURE optimization.**

For biobank-scale executions estimating a high number of ancestral populations ($K > 10$), a subset of hyperparameters is adjusted to maintain stable convergence in a more complex optimization landscape. The modified values for these specific scenarios are provided in Table S6.

| Parameter | Description | Value |
|---|---|---|
| $\alpha$ | Initial learning rate. | 0.0075 |
| $\gamma$ | Decay factor. | 0.85 |
| $t_{\text{max}}$ | Maximum iterations. | 10000 |
| $\rho$ | Learning rate decay patience. | 5 |

**Table S6. Modified hyperparameters for biobank-scale executions with $K > 10$.**

#### SUPPLEMENTARY NOTE S5: COMMON LIBRARIES, ENVIRONMENT AND PARAMETERS

Expanding on the hardware configuration detailed in the experimental setup, benchmarks were performed on an x86_64 Linux system (AMD Zen) using `OpenBLAS v0.3.27` for linear algebra, compiled with `GCC v10.2.1`. To ensure fair comparative benchmarking, memory allocation was standardized across all methods: each process was strictly limited to 50 GB of RAM for standard datasets, which was increased to 256 GB specifically to accommodate the massive scale of the UK Biobank runs.

The core software pipeline was developed using `Python 3.10`. The specific library dependencies and environment requirements for our proposed method, `ADAMIXTURE`, are detailed in Table S7.

Due to significant versioning conflicts between external libraries—specifically the requirement for `NumPy` 2.0 in modern high-performance frameworks versus the legacy `scikit-learn` and `PyTorch` requirements of deep learning baselines—execution was managed through isolated environments. Methods such as `Neural Admixture` and `fastmixture` were executed in dedicated virtual environments to satisfy their mutually exclusive dependency trees, as detailed in Table S8 and Table S9, respectively. In contrast, `ADMIXTURE` and `SCOPE` were maintained as standalone compiled binaries and did not rely on these specific Python package constraints. Finally, to ensure full reproducibility of the comparative analysis, the exact software repositories, and specific execution hyperparameters (including convergence tolerances, iteration limits, and initialization seeds) for all baseline methods are summarized in Table S10.

| Library | Version | Primary Role in the Pipeline |
|---|---|---|
| `NumPy` | $> 2.0.0$ | Fundamental linear algebra and array operations. |
| `Cython` | $> 3.0.0$ | Optimized C-extensions for performance bottlenecks. |
| `PyTorch` | $\geq 2.0.0$ | Tensor operations and hardware (GPU) acceleration. |
| `pgenlib` | $\geq 0.93.0$ | Efficient parsing of PLINK 2.0 binary genotypes (.pgen). |
| `scikit-allel` | $\geq 1.3.8$ | Parsing of VCF files. |
| `Matplotlib` | $\geq 3.5.0$ | Visualization of admixture proportions and convergence. |

**Table S7.** Software environment and library versions for `ADAMIXTURE`.

| Library | Version | Primary Role in the Pipeline |
|---|---|---|
| `NumPy` | 1.21.0– $< 2.0.0$ | Array operations and compatibility with `torch`. |
| `Cython` | $\geq 0.29.30$ | Performance-critical extensions. |
| `PyTorch` | $\geq 1.7.1$ | Autoencoder training and GPU acceleration. |
| `Scikit-learn` | $\leq 1.1.1$ | PCKMeans initialization and PCA preprocessing. |
| `Dask` | 2022.5.0 – 2024.2 | Parallelized data handling and loading. |
| `Pandas-plink` | $\geq 2.2.9$ | Parsing of PLINK binary genotypes. |
| `WandB` | $\geq 0.12.17$ | Experiment tracking and logging. |

**Table S8.** Software environment and library versions for `Neural Admixture`.

| Library | Version | Primary Role in the Pipeline |
|---|---|---|
| `NumPy` | $> 2.0.0$ | Fundamental linear algebra and array operations. |
| `Cython` | $> 3.0.0$ | Optimized C-extensions for EM updates. |

**Table S9.** Software environment and library versions for `fastmixture`.

| Method | Version | Description & Source | Key Parameters |
|---|---|---|---|
| ADMIXTURE | 1.4.1 | An exact EM method utilizing block relaxation and quasi-Newton acceleration. https://dalexander.github.io/admixture/ | Method: block; Acceleration: qn3; $\Delta L$ tol: $10^{-4}$; Max Iter: 1000; Seeds: $\{42, 43, 44, 45, 46\}$. |
| Neural Admixture | 1.6.7 | A deep learning method utilizing an autoencoder architecture. https://github.com/AI-sandbox/neural-admixture | LR: $10^{-5}$; Epochs: 50; Batch size: 400; Init: pckmeans; PCA comps: 2; L2 penalty: $5 \cdot 10^{-4}$; Seeds: $\{42, 43, 44, 45, 46\}$. |
| fastmixture | 1.2.0 | A mini-batch EM method utilizing quasi-Newton acceleration. https://github.com/Rosemeis/fastmixture | Batches: 32; $\Delta L$ tol: $10^{-9}$; Max Iter: 1000; SVD Subsample: 0.7; SVD Power Iter: 11; Seeds: $\{42, 43, 44, 45, 46\}$. |
| SCOPE | Default | A likelihood-free method utilizing alternating least squares. https://github.com/sriramlab/SCOPE | Max Iter: 1000; Convergence tol: $10^{-5}$; Seeds: $\{42, 43, 44, 45, 46\}$. |

**Table S10. Software versions, source URLs, and default execution parameters for baseline methods.**

### SUPPLEMENTARY NOTE S6: QUANTITATIVE CLUSTER STABILITY ACROSS RUNS

To evaluate the robustness and reliability of the inferred ancestral components, we assessed the stability of the admixture proportions ($Q$) across independent runs with different random seeds. Since the indexing of clusters (e.g., which population is labeled as "Cluster 1") can vary between executions, we first perform a greedy alignment of the $Q$-matrices by minimizing the squared Euclidean distance between their columns.

The stability is quantified using two pairwise metrics averaged across all combinations of runs. For any two aligned admixture matrices $Q^{(a)}$ and $Q^{(b)}$ of size $N \times K$ (where $N$ is the number of individuals and $K$ is the number of ancestral populations), we define:

1. **Mean Pearson Correlation:** Measures the linear consistency of the ancestral assignments across runs. It is calculated as the average column-wise correlation:

$$\rho(Q^{(a)}, Q^{(b)}) = \frac{1}{K} \sum_{k=1}^{K} \frac{\sum_{i=1}^{N} (q_{ik}^{(a)} - \bar{q}_k^{(a)})(q_{ik}^{(b)} - \bar{q}_k^{(b)})}{\sqrt{\sum_{i=1}^{N} (q_{ik}^{(a)} - \bar{q}_k^{(a)})^2} \sqrt{\sum_{i=1}^{N} (q_{ik}^{(b)} - \bar{q}_k^{(b)})^2}} \tag{3}$$

   where $\bar{q}_k$ represents the mean ancestral proportion for cluster $k$. Values near 1.0 indicate high replicability.

2. **Frobenius Distance:** Measures the overall divergence between aligned $Q$-matrices, defined as the Frobenius norm of their difference:

$$d_F(Q^{(a)}, Q^{(b)}) = \|Q^{(a)} - Q^{(b)}\|_F = \sqrt{\sum_{i=1}^{N} \sum_{k=1}^{K} \left( q_{ik}^{(a)} - q_{ik}^{(b)} \right)^2} \tag{4}$$

   A lower distance indicates that the optimization consistently converges to the same region of the global likelihood surface.

As shown in Figure S1, ADAMIXTURE achieves near-perfect stability (Correlation $> 0.99$) and minimal Frobenius distances across most datasets, outperforming or matching existing baselines while significantly reducing the variance between initializations.

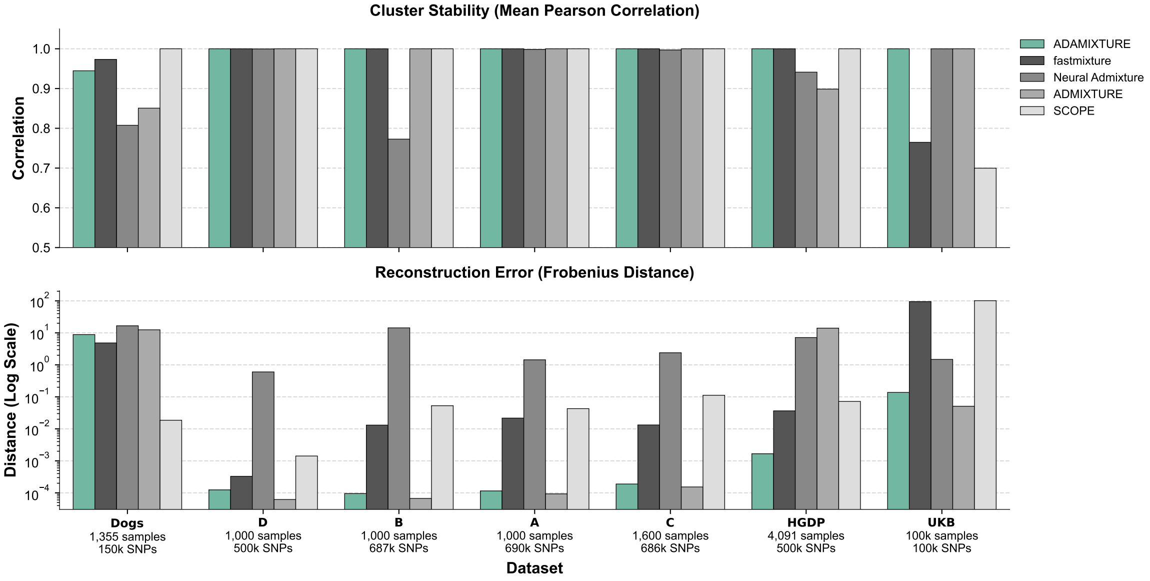


Figure S1: **Quantitative Cluster Stability.** Top panel: Mean Pearson Correlation between aligned $Q$-matrices from independent runs. Bottom panel: Pairwise Frobenius Distance (log scale) between aligned runs. Metrics were calculated across 5 independent seeds for each method and dataset.

### SUPPLEMENTARY NOTE S7: FIGURE ACCESSIBILITY DESCRIPTIONS

To comply with the Oxford University Press (OUP) accessibility policy and the ISMB 2026 guidelines, the following alternative text descriptions are provided for the figures located in the main manuscript:

- **Figure 1 Alt-Text:** A five-panel stacked bar chart comparing individual ancestry proportions inferred by ADAMIXTURE, fastmixture, Neural Admixture, ADMIXTURE, and SCOPE across six geographic regions (England, India, Wales, Scotland, Republic of Ireland, and Northern Ireland). ADAMIXTURE, fastmixture, and classical ADMIXTURE display visually indistinguishable, well-defined genetic clusters. Conversely, Neural Admixture displays severe over-smoothing with a single dominant cluster across all regions, and SCOPE exhibits noisy, poorly resolved boundaries, visually confirming the superior accuracy of the EM-based methods.

- **Figure 2 Alt-Text:** A grouped bar chart on a logarithmic y-axis comparing the execution runtimes of six methods across seven datasets of varying sizes. The classic ADMIXTURE method consistently exhibits the tallest bars, approaching 100 hours on the largest dataset (UKB). In stark contrast, both ADAMIXTURE (GPU) and ADAMIXTURE (CPU) consistently display the lowest bars alongside SCOPE, completing most tasks in under a few minutes and visually demonstrating an orders-of-magnitude speedup over classical ADMIXTURE and fastmixture.

- **Figure 3 Alt-Text:** A line plot on a logarithmic y-axis illustrating runtime scalability as a function of the number of ancestral populations (K) on the x-axis, ranging from 5 to 50. The line for fastmixture rises steeply, indicating rapid exponential growth in execution time, and terminates abruptly at K=20. In contrast, the lines for ADAMIXTURE (CPU) and ADAMIXTURE (GPU) demonstrate significantly flatter, near-logarithmic trajectories across the entire range up to K=50, with the GPU version consistently maintaining the lowest runtime, barely exceeding 1 hour even at maximum complexity.

- **Figure 4 Alt-Text:** A four-panel line graph plotted on a logarithmic y-axis, illustrating the execution time scaling of ADAMIXTURE (GPU), ADAMIXTURE (CPU), and fastmixture. The top panels show time versus the number of SNPs (from 100k to 500k) for fixed sample sizes of 100,000 and 500,000. The bottom panels show time versus the number of samples (from 50k to 500k) for fixed SNP counts of 100,000 and 500,000. Across all scenarios, fastmixture exhibits the steepest growth, approaching 100 hours at maximum dimensions. ADAMIXTURE (CPU) shows improved scalability, while ADAMIXTURE (GPU) consistently displays the lowest and flattest trajectories, maintaining execution times under two hours even at the half-million scale for both samples and SNPs.
